# Dipeptidyl dipeptidase-4 inhibitor, MK-0626, promotes bone marrow-derived endothelial progenitor cell bioactivities for vascular regeneration in diet-induced obese mice

**DOI:** 10.1101/429399

**Authors:** Amankeldi A. Salybekov, Haruchika Masuda, Kozo Miyazaki, Yin Sheng, Atsuko Sato, Tomoko Shizuno, Yumi Iida, Yoshinori Okada, Takayuki Asahara

**Affiliations:** Department of Regenerative Medical Science, Division of Basic Clinical Science, Tokai University School of Medicine, Isehara, Japan; The Support Center for Medical Research and Education, Tokai University School of Medicine, Isehara, Japan

**Author notes:** Corresponding author Takayuki Asahara, MD, Ph.D., Department of Regenerative Medicine Science, Tokai University School of Medicine, 143 Shimokasuya, Isehara, Kanagawa 259–1193, Japan.

## Abstract

Metabolic syndrome (MS), overlapping type 2 diabetes, hyperlipidemia, and/or hypertension, based on high-fat diet, poses risk for cardiovascular disease. A critical feature associated with such risk is the functional impairment of endothelial progenitor cells (EPCs). Dipeptidyl dipeptidase-4 inhibitors are known not only to inhibit degradation of incretins to control blood glucose levels, but also to improve EPC bioactivity and induce anti-inflammatory effects in tissues. In the present study, we investigated the effects of such an inhibitor, MK-06266, in ischemia model of MS using diet-induced obese (DIO) mice. EPC bioactivity was examined in MK-0626-administered DIO mice and non-treated control group, using an EPC colony-forming assay and bone marrow cKit^+^ Sca-1^+^ lineage-cells, and peripheral blood-mononuclear cells. Our results showed that, *in vitro*, the effect of MK-0626 treatment on EPC bioactivities and differentiation was superior in comparison with non-treatment. Further, *in vivo* hindlimb ischemia model experiment indicated that microvascular density and pericyte-recruited arteriole number were increased in MK-0626-administered group, but not control group. Lineage profiling of isolated cells from ischemic tissues disclosed that MK-0626 administration has an inhibitory effect on unproductive inflammation. This occurred via a decrease in the influx of total blood cells and pro-inflammatory cells such as neutrophils, total macrophages, M1, total T-cells, cytotoxic T-cells, and B-cells, with a concomitant increase in number of regeneration-associated cells, such as M2/M ratio and T_reg_/T-helper. Laser Doppler analysis revealed that at day 14 after ischemic injury, blood perfusion in hindlimb was grater in DIO mice treated with MK-0626, but not in control. In conclusion, the dipeptidyl dipeptidase-4 inhibitor has a positive effect on EPC differentiation in MS model of DIO mice. Following ischemic injury, DPP-4 i sharply reduces recruitment of pro-inflammatory cells into ischemic tissue, and triggers regeneration and reparation process. Thus, DPP-4 i is a promising therapeutic agent for MS treatment.

## Introduction

Vascular regeneration is an initial and essential process for organ regeneration. Endothelial progenitor cells (EPCs) play key roles in vasculogenesis [1] and in regulating angiogenesis [2]. Thus, EPC kinetics and bioactivities are essential for vascular regeneration and organogenesis in regenerative medicine. Type 2 diabetes is associated with the decrease in number and impaired function of circulating EPCs, which in turn has been linked to cardiovascular disease complication [3]. There are several options to stimulate EPC proliferation and biological functions in diabetes that are being actively pursued by researchers [4]. For example, several cytokines such as stromal cell-derived factor-1 (SDF-1), granulocyte colony stimulating factor (G-CSF), and granulocyte macrophage colony stimulating factor (GM-CSF), angiogenic growth factors such as vascular endothelial growth factor (VEGF), and pharmaceutical drugs such as estrogen and statins, have all been reported to augment EPC bioactivities, such as proliferation, differentiation, migration, mobilization, and recruitment of BM-derived EPCs [5, 6].

Dipeptidyl dipeptidase-4 (DPP-4) inhibitors (DPP-4 i) have been broadly applied in clinical aspects for controlling blood glucose levels in type 2 diabetic patients (T2DP). DPPIV i inhibits the degradation of incretins, such as glucagon-like peptide-1 (GLP-1), glucose-dependent insulinotropic polypeptide (GIP), leading to increased insulin secretion from Langerhans islets [7]. Notably, DPP-4 also targets other physiological substrates, especially functional cytokines regulating stem/progenitor bioactivities, e.g., stromal derived factor-1α (SDF-1α) [8]. Further, studies have shown that DPP-4 is critical for the mobilization of EPCs from the bone marrow [9].

Fadini et al. demonstrated that the DPP-4 i sitagliptin increases the level of circulating EPCs in type 2 diabetic patients, with concomitant up-regulation of SDF-1 [10]. However, at that time, EPC research methodologies were still relatively underdeveloped to obtain clear scientific insights into stem cell biology for vascular regeneration. Recently, our laboratory developed new EPC research methodologies, EPC-CFA (new EPC colony forming assay) and QQ-EPC culture (quality and quantity controlled serum-free EPC culture technique), to identify a variety of EPC phenotypes and functions[11]. These methodologies allow us to precisely evaluate the key factors of EPC kinetics and bioactivities, and ultimately lead us to understand how EPC differentiates in healthy or diseased states to promote new vasculature.

In this study, we investigated whether a sitagliptin analogue, MK-0626, affects EPC kinetics in peripheral blood or bone marrow in diabetic animal models, using the new EPC quality and quantity evaluation methods. Furthermore, the developed cell isolation technique from ischemic muscles was used to define and compare in situ cell phenotype and quantity of hematopoietic cells in tissues, in order to evaluate inflammatory cell kinetics in diabetic animals following MK-0626 administration.

## Materials and methods

All studies were performed with the approval of national and institutional ethics committee. The Tokai School of Medicine Animal Care and Use Committee gave local approval for these studies, based on Guide for the Care and Use of Laboratory Animals (National Research Council).

### Reagents

Dipeptidyl dipeptidase-IV (DPP-4) inhibitor (DPP-4 i) MK-0626 was gifted by MSD K. K. (Kenilworth, N.J., USA.). MK-0626 was dissolved in 0.25% methyl cellulose (M-0389, SIGMA, St. Louis, MO, USA) solution for *per os* administration [12].

### Animals

Ten-to fifteen-week-old C57BL/6J (Lean) and C57BL/6J DIO (DIO) male mice were purchased from Charles River Laboratories (Yokohama, JAPAN) via Oriental Yeast Co. Ltd. (Tokyo, JAPAN) and maintained under the standard conditions (20±2°C, relative humidity (50–60%), light/dark 12h/12hcycles) at the Support Center for Medical Research and Education in Tokai University, School of Medicine.

During one week of acclimatization, C57BL/6J mice received a standard rodent diet, in which 10% of energy came from fat (D12450J, Research Diet Inc., New Brunswick, NJ, USA), while C57BL/6J-DIO mice received a high fat diet (HFD), in which 60% energy came from fat (D12492, Research Diet Inc., New Brunswick, NJ, USA). After three weeks of feeding with the respective diets, mice were divided into two groups. The solution of MK-0626 was daily administered to mice of each group by oral cannulation with sonde (3 mg/kg/day) for 1 week. Based on previous report, this dose of MK-0626 was predicted to result in continuous blocking of incretins, such as GLP-1 and GIP, and inactivation of DPP-4[13]. Food intake of the mice was recorded every two days and their body weight (BW) and blood sugar (BS) were measured 9 and 3 days before surgery, and on day 4 and day 11 after surgery. Based on BW at each time point, the volume of MK-0626 solution was adjusted to maintain the same dose in each mouse until the subsequent measurement. The BS was measured using a blood glucose test meter (Gultest Ace R, ARKRAY Factory, Inc. Shiga, Japan) and disposable blood glucose level test sensor (Gultest sensor, Panasonic Healthcare Co., Ltd.).

At the end of the experimental period, the mice were anesthetized with pentobarbital and their plasma collected and stored at –80°C.

### Cell preparation and culture

Mouse peripheral blood mononuclear cells (mPBMNCs) were collected with 27G-Insulin syringe (TERUMO, Tokyo, Japan) from the apex of the heart under adequate anesthesia using 1.5% to 2.0% isoflurane (Dainippon Sumitomo Pharma Co., Ltd., Osaka, JAPAN). The cells were further isolated by density gradient centrifugation with Histopaque (d=1.083; Sigma-Aldrich Co., St Louis, MO, USA), as previously reported[14].

Preparation of human peripheral blood mononuclear cells (hPBMNCs) was performed after obtaining informed consent from healthy volunteers according the Tokai University institutional the review board. The peripheral blood (10 mL) was drawn by heparinized venous puncture at the forearm. Isolation protocol for hPBMNCs was the same as that for mPBMNCs. Briefly, cells were cultured with QQ (Quality and Quantity) culture media of Stem Line II (Sigma-Aldrich, St. Louis, MO), supplemented with 100 ng/mL recombinant mouse (rm) or human (rh) stem cell factor (SCF), 100 ng/mL Flt-3 ligand (FL3L), 20 ng/mL thrombopoietin (TPO), 50 ng/mL vascular endothelial growth factor (VEGF), 20 ng/mL interleukin-6 (IL-6) (these five proteins were purchased from Peprotech, Inc. (Rocky Hill, NJ, USA), and antibiotics Penicillin/Streptomycin (100 U/100 µg/mL; Gibco). The cells (5 × 10^5^ cells) were cultured for 3 days (mouse mPBMNCs) or 7 days (human hPBMNCs) MNCs on 24-well plate (BD Falcon, BD Bioscience, San Jose, CA, USA) in a 37°C incubator containing humidified atmosphere with 5% CO_2_.

### Enrichment of EPCs from bone marrow

Bone marrow (BM) EPCs were isolated from no-fat diet and DIO-mice femurs and tibias as previously described [15]. Nuclear cells were washed with PBS-EDTA and the erythrocytes were removed by hemolyzation lysis buffer. Nuclear cells were initially stained with a lineage positive antibody cocktail containing CD45R/B220, TER119, CD3e, CD11b, Ly-6G, and Ly6C (Gr-1) for 20 minutes at 4°C (all antibodies were obtained from BD Pharmingen). After labeling the lineage positive antibodies with biotin-labeled magnetic beads, cells underwent a negative selection process with a magnetic cell sorting system (Auto MACS^™^, Miltenyi Biotec.). An EPC-enriched BM population (Lin^-^, c-kit^+^, Sca-1^+^; KSL) was isolated by FACS Aria^™^ cell sorter (BD) from BM-Lin^-^cells. The Lin^-^ cells were counted and then incubated with Rat-FITC anti-mouse Ly-6A/E (Sca-1) (BD PharMingen) and Rat-PE CD117 (c-kit) (BD PharMingen) for 20 min at 4°C, washed three times and suspended in 20% IMDM (Gibco). FITC-conjugated Sca-1^+^ and PE-conjugated c-Kit^+^ double positive cells (KSL) were obtained using the FACS Aria cell sorter (BD).

### EPC colony forming assay

Freshly isolated human or mouse peripheral blood mononuclear cells (PBMNCs) and mouse BM mononuclear cells (BMMNCs) were cultured in semisolid methyl cellulose-based culture medium, (MethoCult^™^ SF M3236, STEMCELL Technologies Inc., Vancouver, BC, Canada) containing 100 ng/mL SCF, 50 ng/mL VEGF, 50 ng/mL basic fibroblast growth factor (bFGF), 50 ng/mL epidermal growth factor (EGF), 50 ng/mL insulin-like growth factor (IGF), 50 ng/mL interleukin-3 (IL-3) (these six proteins were purchased from Peprotech, Inc. Rocky Hill, NJ, USA), 2 IU/mL heparin (Ajinomoto Pharmaceutical Co. Ltd. Tokyo, Japan), 30% (v/v) fetal bovine serum (Nichirei Biosciences Inc., Tokyo, Japan) and penicillin/streptomycin (100 U/100 µg/mL). Cells were seeded at 1.5 × 10^5^ cells/35 mm dish (BD Falcon, BD Bioscience, San Jose, CA, USA) and left in a humidified incubator with 5% CO_2_ at 37°C till EPC colony formation. The number of adherent colonies on the dishes was counted between day 6–10 (mouse) and 16–18 (human) using gridded scoring dish (STEMCELL Technologies Inc. Vancouver, BC, Canada) under a phase-contrast light microscope (Eclipse TE3000; Nikon, Tokyo, Japan). Primitive EPC colony-forming units (pEPC-CFUs) and definitive EPC colony-forming units (dEPC-CFUs) were separately counted.

### Endothelial lineage characterization

As previously described [16], after culturing PBMNCs for 7 days in the endothelial cell growth medium (EGM-2 MV BulletKit, Lonza, Walkersville, MD, USA), we evaluated enrichment of the EPC lineage by staining with Fluorescein Ulex Europaeus Agglutinin I (UEA-I Lectine, FL-1061, Vector Laboratories Inc. Burlingame, CA, USA) and acetylated low-density lipoprotein labeled with 1,1’-dioctadecyl-3,3,3’,3’-tetramethylindo-carbocyanine perchlorate (DiI-Ac LDL, BT-902, Biomedical Technologies Inc. St. Stoughton, MA, USA), and then observed the cells under the Bio Revo fluorescence microscope (BZ-9000, Keyence, Osaka, Japan). EGM-2-MV complete medium was adjusted to EBM-2 basal medium by adding 5% FBS (SAFC Biosciences Inc., Lenexa, KS) and supplemented with growth factors, except hydrocortisone. PBMNCs were adjusted to the similar cell density (1 × 10^6^ cells/mL) with EGM-2-MV complete medium containing 5% FBS. Cells were then plated on 6-well Primaria tissue culture plate (2 × 10^6^ cells/2 mL per well) and cultured.

### Induction of hindLimb ischemia model

The mice were divided into two groups, Lean and DIO, one week before surgery. Hindlimb ischemia induction (HLI) was performed under adequate anesthesia by 1.5% to 2% isoflurane to minimize pain, according to the 3Rs rule (replacement, reduction, and refinement). Briefly, the proximal portion of the left common and deep femoral artery with their three branches were successfully ligated with a 6–0 nylon suture (Sigma Rex., Kono manufacturing Co., Ltd. Ichikawa, Japan), and the proximal and distal portions of the saphenous artery were subjected to bipolar electrocautery (MERA N3–14 SENKO MEDICAL INSTRUMENT mfg. Co., Ltd., Tokyo, Japan). The skin was closed with a 4–0 nylon suture (Sigma Rex., Kono manufacturing Co., Ltd. Ichikawa, Japan). To reduce post-surgery suffering from pain, pentobarbital (Kyouritu Seiyaku Co., Ltd., Tokyo, Japan) was injected at a dose of 10 mg/kg via i.p. administration.

### Laser doppler imaging and blood flow assessment

Baseline laser Doppler perfusion imaging (LDPI; Moor Instrument, Axminster, UK) was performed of animals under anesthesia with 1.5% isoflurane (Dainippon Sumitomo Pharma) and after induction of ischemia at days 0, 7, and 15 to assess blood perfusion ratio in ischemic vs. healthy hindlimb. Acquired data using LDPI were analyzed with moorLDI^™^ Main software (Laser Doppler Imager ver 5.2; Moor Instruments, Devon, UK). Mice with toe necrosis or limb salvage were included in the study, whereas those with foot necrosis or autoamputation were excluded.

### Identification of phenotypes of recruited cells isolated from ischemic tissues

All mice were fed with 5 g HFD per day till enough BS and BW could be retained; then they were divided into two groups, control (received 0.25% methylcellulose only) and MK-0626 group (MK-0626 was administrated *per os* using sonde 3 days before and 3 days after onset of LHI). At day seven, mouse was sacrificed after anesthesia, and systemically perfused with cold PBS to exclude blood cells. An anterior tibial muscle (ATM) was dissected for further isolation of cells that had infiltrated the ischemic tissue. Our previous immunohistochemistry analysis study showed that ATM is the most sensitive for ischemic injury. In brief, ATM muscle vessels, tendons and nerve fibers were removed under light microscope and minced by using optical fine micro scissors. To effectively liberate skeletal muscle cell types, collagenase type II (500 U/mL) (Worthington Lab.) and collagenase/dispase (1 mg/mL) (Roshe Diagnostics) were used for 1.5 h at 37°C with gentle agitation, as reported elsewhere [17]. After digestion, the tissue was triturated and meshed through a 70-µm cell strainer. Finally, cells were washed twice with DMEM (Gibco) and then counted using a hemocytometer. The Fcλ receptors were blocked with mouse anti-Fcλ receptor (Biolegend Co. Ltd. CA, USA) to reduce nonspecific binding of antibodies and left at 4°C for 30 min and then washed twice with FACS buffer. Subsequently, cells were stained with the mixture of antibodies (Biolegend Co. Ltd. CA. USA) against CD45, CD34, CD206, F4/80, CD11b, Ly-6G, CD31, Sca-1, CD117, CD3e, CD4, CD8a, CD25 and CD19 at 4°C for 40 min after which the cells were washed twice. Flow cytometric analysis was performed on a BD FACS Verse and Fortessa (BD), and data was analyzed using FlowJo (TreeStar 10.2 version) and the DeNova software version 6.

### Immunohistochemistry analysis

Two weeks after surgery, the mice were injected with 20 µL of fluorescein isothiocyanate-conjugated *Griffonia simplicifolia* isolectin B4 (IB4-FITC, FL1201, Vector Laboratories Inc.) via tail vein to detect functional capillaries *in vivo*. Twenty minutes after injection, the mice were sacrificed under overdose of pentobarbital 150 mg/kg/ml (via i.p administration), and then systemically perfused animals were fixed with 4% paraformaldehyde. Ischemic tissues were left in 4% paraformaldehyde overnight at 4°C, and anterior tibial muscles were excised, and embedded into paraffin block. The deparaffinized tissue sections were mounted on slide glass with glycerol-PBS solution (pH 8.0) including DABCO (D27802–25G SIGMA-Aldrich) for preventing photobleaching. To evaluate pericyte recruitment, Cy3-conjugated monoclonal anti-actin alpha-smooth muscle (α–SMA) antibody (1:200, clone: 1A4, Sigma-Aldrich), and for micro vascular density (MVD) IB4-FITC (1:1000, Invitrogen) was used. The tissue sections for MVD and pericyte recruitment were observed and counted using fluorescence microscopy VH Analyzer (Keyence).

### Statistical analysis

All values are shown as mean ± SE. Mann-Whitney U and Kruskal-Wallis test were used for two and three non-parametric groups with Dunn’s multiple comparison test, respectively. For multiple comparisons between groups at different time points, 2-way ANOVA was applied, followed by Tukey’s post hoc test. All statistical analyses were performed using GraphPad Prism 7.1 (GraphPad Prism Software Inc., San Diego, CA, USA). *P* < 0.05 value was considered to indicate statistically significant differences.

## Results

### EPC differentiation was induced by MK-0626

To evaluate the efficacy of MK-0626 on EPC’s colony forming ability and differentiation, PBMNCs were isolated from Lean mice, MK-0626-administered Lean mice, DIO mice, and MK-0626-administered DIO mice, and cultured in semisolid culture media. The number of definitive EPC-CFUs (dEPC-CFUs) from control DIO mice decreased compared to that from control Lean mice, whereas dEPC-CFUs from MK-0626-administered DIO mice were similar in number to those from the Lean littermates (Fig 1A). The frequency of dEPC-CFUs per peripheral blood (PB) volume also sharply decreased in DIO mice compared with that in control mice, while MK-0626-administered mice showed a dramatic improvement in the number of dEPC-CFUs per 1 mL PB (Fig 1B). These data suggest that EPC differentiation in PBMNCs was impaired in DIO mice and MK-0626 treatment effectively recovered the EPC differentiation ability to similar levels seen in Lean mice. We further investigated whether DPP-4 i administration affected bone marrow c-Kit^+^/Sca-1^+^/Lin^-^ (BM-KSL) stem cells in obese and healthy condition. To address this, purified BM-KSL stem cells, isolated from either control (lean) or DIO mice, were evaluated for their EPC colony formation abilities. As shown in Fig. 1C, BM-KSL cells displayed similar EPC-CFUs between all tested groups, indicating that the EPC differentiation ability was not affected by obese condition or MK-0626 administration in BM. However, the calculated number of EPC-CFUs in hemi bone represented stimulated expansion of EPCs in DIO mice, and MK-0626 did not affect the frequency of EPC-CFUs in BM in both DIO and Lean mice (Fig 1D).

**Fig 1.**
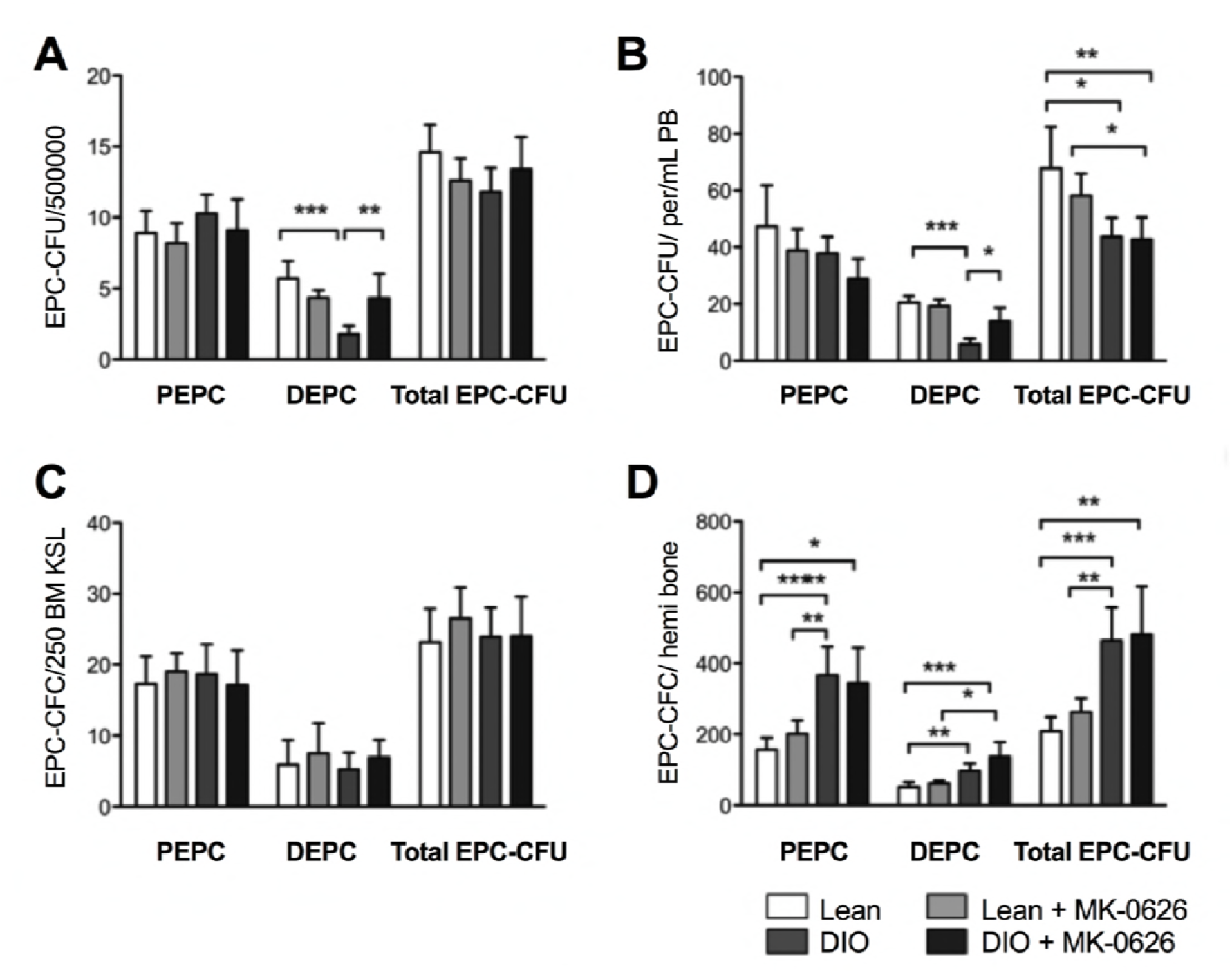
MK-0626 enhances EPC differentiation *in vivo*. The colony number of pEPC-CFUs and dEPC-CFUs generated from 5 × 10^5^ cells of BM (A), 1 mL of peripheral blood (B), 250 cells of BM-KSL (C) and hemi bone (D). The individual bars indicate cells from healthy mice (open), MK-0626-administered healthy mice (light gray), DIO mice (dark gray) and MK-0626-administered DIO mice (black). Data are represented as the mean ± SE. N = 6 mice per group. Experiments were repeated twice. In the graph, \**P* < 0.05, \*\**P* < 0.01,\*\*\**P* < 0.001, and \*\*\**P* < 0.0001 as determined by Two-way ANOVA followed Tukey’s multiple comparisons test.

To investigate MK-0626 effect on EPC biology, we employed ex-vivo regenerative conditioning in culture with or without MK-0626 on BM-derived KSL cells from DIO or Lean mice. After one week of culturing, colony-forming assay (CFA) was used to evaluate these cells. Conditioning BM-KSL cells with MK-0626 promoted the EPC colony forming potential compared with Lean mice without MK-0626 treatment (Fig 2B). However, EPC-CFA values were higher in DIO mice compared with those in control lean mice, and did not further improve upon MK-0626 administration. Two thirds of the EPCs in treated and non-treated Lean mice were in the pEPC stage, while half of the EPCs in treated and un-treated DIO mice were definitive EPC colonies, which are responsible for vasculogenesis (Fig 2C).

**Fig 2.**
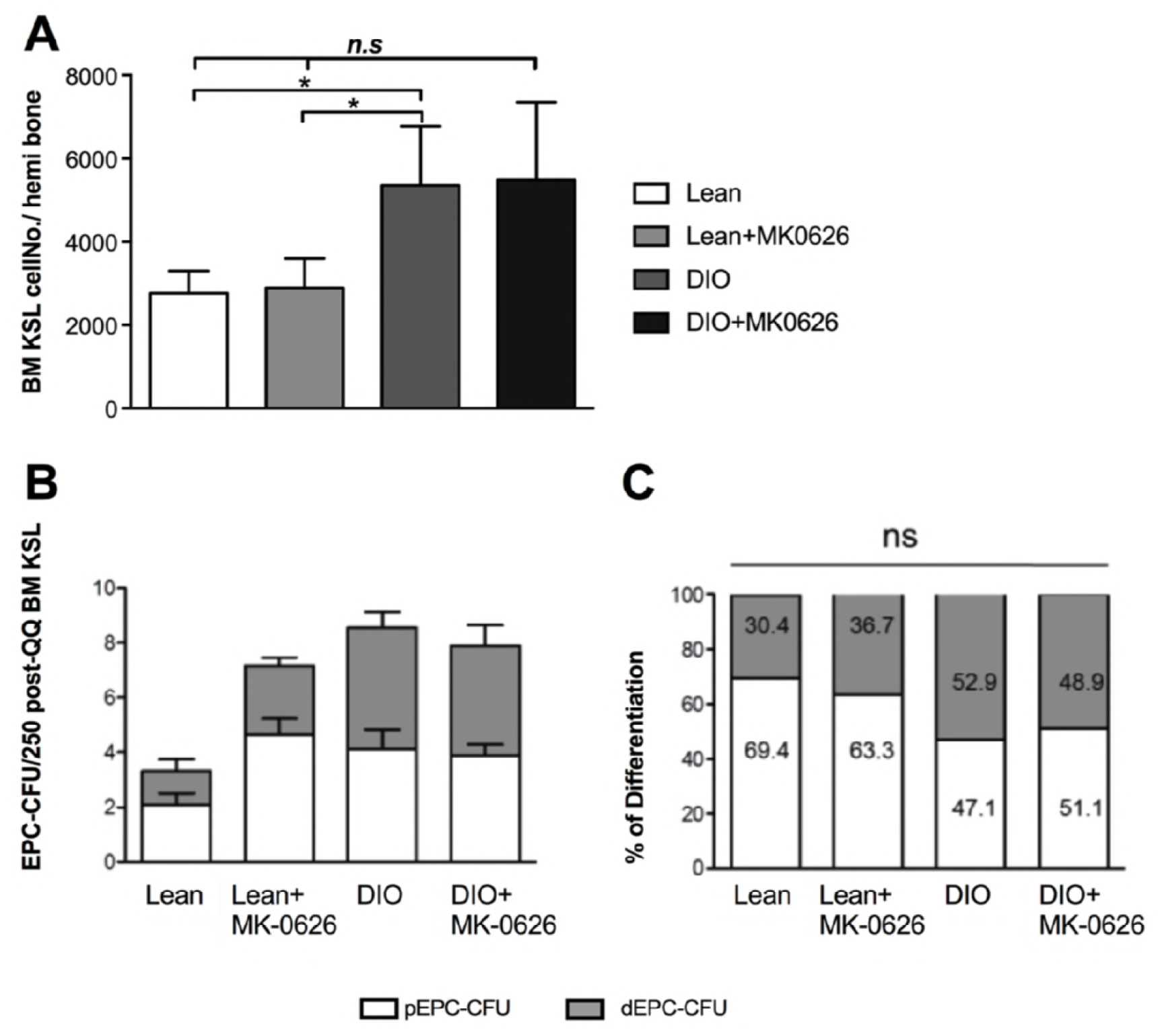
Differentiation and mobilization ability of BM-KSL cells treated with MK-0626. (A) The graph shows counts of BM-KSL cells isolated from a pair of femur and tibia at a hemi-side in each mouse. (B) EPC-CFU counts generated from BM-KSL cells per dish (250 cells/dish), and also shows the percentage of differentiation of pEPC-CFU count versus dEPC-CFU per dish (C). The bars indicate the counts of pEPC-CFU (open) and dEPC-CFU (light gray). Data are represented as mean ± SE. N = 6 mice per group. Experiments were repeated twice. In the graph, \**P*< 0.05, \*\**P*<0.01, and \*\*\**P* < 0.001, as determined by Kruskal-Wallis and Dun’s multiple comparison test.

### MK-0626 recovered human EPC colony formation and differentiation capability under inflammatory conditions

To verify the effect of MK-0626 on human EPCs, we performed EPC-CFA and EPC culture assay on regenerative conditioned PBMNC. Diet-induced obesity (DIO) develops systemic chronic inflammatory milieu, by increasing secretion of pro-inflammatory cytokines, such as TNFα, IL-6 and IL-1b[18, 19]. Based on this, we attempted to assess whether EPC differentiation cascade differs under a normal regenerative condition (QQ culture) or an inflammatory condition induced by TNFα (QQ + TNFα). EPC differentiation was evaluated by EPC-CFA before and after QQ culture of hPBMNCs. Inflammatory conditioning markedly decreased EPC colony forming units (Fig 3Ac and 3B), suggesting that inflammatory conditioning significantly impairs EPC colony formation bioactivity. In contrast, MK-0626 supplementation together with TNFα beneficially recovered the EPC function under inflammatory conditioning (Fig 3Ad and 3B), while MK-0626 treatment alone did not increase EPC activity in regenerative conditioning. This may suggest that under regenerative conditioning healthy EPC activation is at its peak and may not respond to additional stimulation with MK-0626. Furthermore, co-staining showed that UEA-I Lectine and DiI-Ac LDL EPCs were decreased significantly in TNFα-treated DIO cells, while combination of MK-0626 and TNFαrecovered the EPC numbers to the same extent as in the healthy Lean group (Fig 3C a through d, and 3D), suggesting that MK-0626 recover EPC function under inflammatory conditions. Together, these results depicted that MK-0626-treatment under inflammatory conditioning favorably enhanced EPC functions such as colony formation and differentiation.

**Fig 3.**
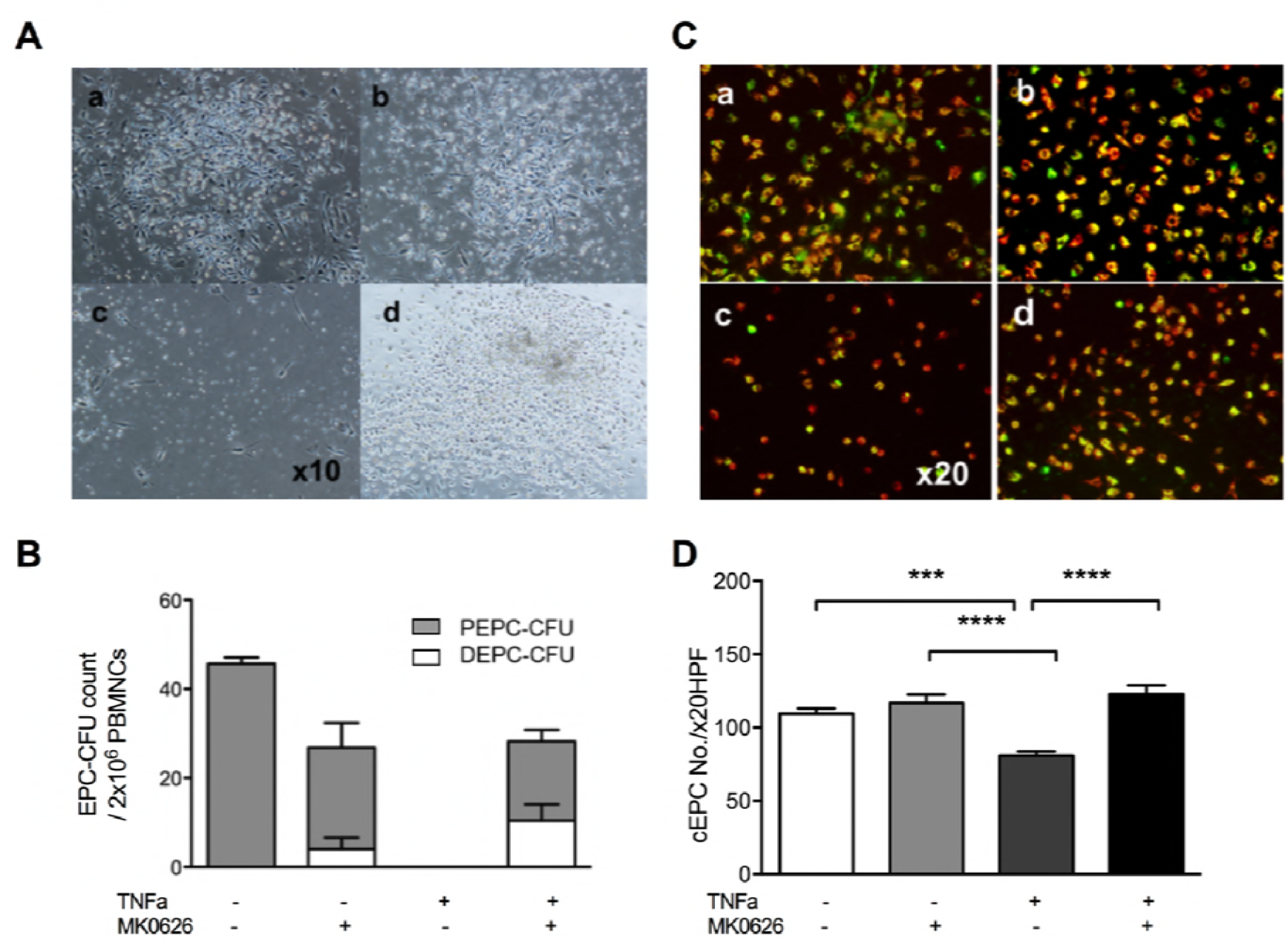
MK-0626 accelerates EPC differentiation under inflammatory conditions. (A) Representative picture of EPC-CFU generated from hPBMNCs. The images are of following EPCs: control (Aa), MK-0626-treated (Ab), TNFα-treated (Ac) and MK-0626- and TNFα-treated (Ad). These EPC-CFUs were observed under high-power field (HPF) of 10×. (B) The graph shows EPC-CFU counts, generated from hPBMNCs (2 × 10^5^ cells/dish). The bars indicate the values of pEPC-CFU (open) and dEPC-CFU (light gray). (C) Conditioned EPCs (cEPCs) were observed with fluorescence microscope after co-staining with UEA-I Lectine (green) for detection of endothelial cell and DiI-Ac LDL (red) for detection of endothelial and macrophage cell lineages. The images are of following cEPCs: control (Ca), MK-0626-treated (Cb), TNFα-treated (Cc), and MK-0626- and TNFα-treated (Cd). The number of the cEPC was counted under high-power field (HPF) of 20×. Data are represented as mean ± SE. N = 3 volunteers. Experiments were repeated at least three times. In the graph, \**P* < 0.05, \*\**P <*0.01, and \*\*\**P* < 0.001, determined by Two-way ANOVA followed by Tukey’s multiple comparisons test.

### MK-0626 administration promoted blood flow perfusion after HLI in DIO mice

To demonstrate in vivo blood flow perfusion recovery, we induced mouse HLI model in healthy and DIO mice and examined the effects using laser Doppler perfusion imaging. After HLI surgery (left-side limb), DIO mice blood flow perfusion deteriorated sharply (0.23 ± 0.03), in comparison with MK-0626-treated DIO mice. Moreover, at day 13 after HLI surgery, DIO mice treated with MK-0626 showed significant improvement of blood flow (0.46 ± 0.03, P <0.05), similar to that seen in the Lean groups (0.47 ± 0.06 and 0.53 ± 0.06 in Lean and Lean treated with MK-0626, respectively) (Fig 4A and 4B). This data showed that DPP-4 i accelerates angiogenesis for further enhancement of blood flow and limb salvage in DIO-conditioned mice.

**Fig 4.**
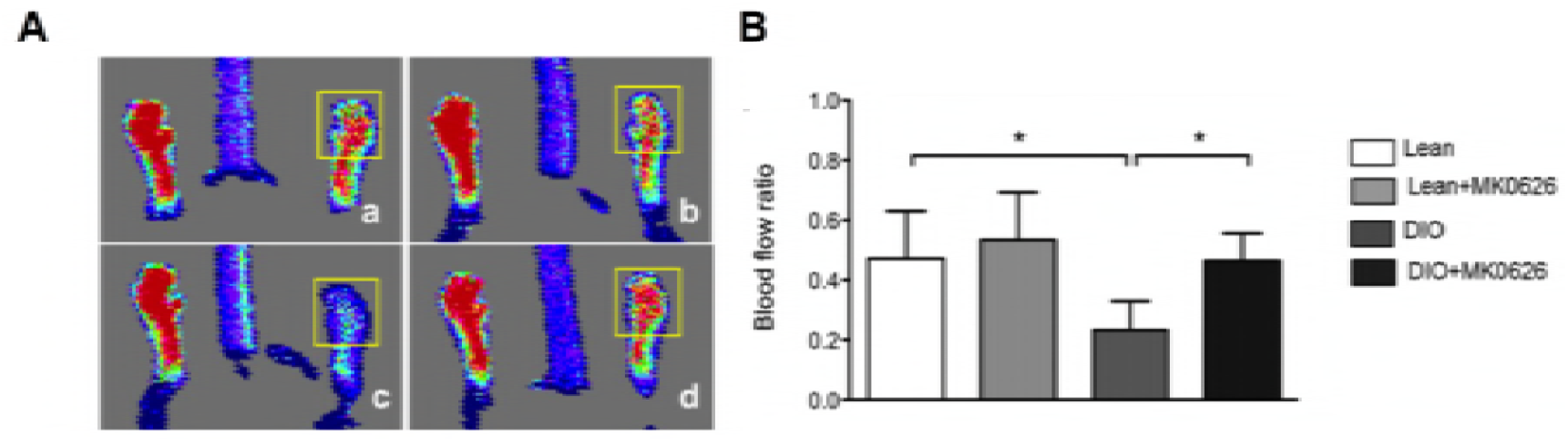
Blood flow recovery rate after HLI. (A) Laser Doppler imaging was used to analyze blood flow 14 days after ischemia. The panels show lean mice (Aa), MK-0626-administered lean mice (Ab), DIO mice (Ac), and MK-0626-administered DIO mice (Ad). The region of interest (ROI) for blood flow measurement is shown by the yellow square. (B) The graph presents blood flow ratio of ischemic-to-contralateral hind limb, during the observation period at day 14. Experiments were repeated twice. In the graph, \**P* < 0.05, \*\**P* < 0.01, and \*\*\**P* < 0.001, determined by Kruskal-Wallis and followed Dun’s multiple comparison test.

### MK-0626 administration reduced infiltration of pro-inflammatory hematopoietic cells lineages

At day 3 after onset of HLI, infiltrated blood cells in the ischemic anterior tibial muscle were successfully isolated and analyzed using flow cytometry for cell subset determination (Fig 5). Viable cell population was gated into two main cell populations, CD45 positive (CD45^+^) or negative (CD45^-^) cells. This strategy led us to determine origin of infiltrated cells, such as blood cells, resident cells or transitional phase cells. HLI surgery caused abundant recruitment of CD45^+^ cells into the ischemic tissue in DIO group, which diminished significantly (5 times less, *P*< 0.028) by MK-0626-treatment (Fig 6A). To determine the inflammation-and regeneration-associated cell proportion among blood cells, we separately gated total macrophages (F4/80^+^), neutrophils (Ly-6G), T-cells (CD3e^+^) and B-cells (CD19^+^) lineages (Fig 5). Numerically, F4/80^+^ and Ly-6G cell accumulations in the ischemic skeletal muscle were significantly increased in DIO mice (*P* < 0.01 and P = 0.057, respectively), in comparison with the MK-0626-treated DIO group (Fig 6B). To evaluate the total macrophage sub-population, we gated two different functional macrophages, pro-inflammatory M1 (CD45^+^/F4/80^+^/CD11b^+^) and anti-inflammatory M2 (CD45^+^/F4/80^+^/CD206^+^) (Fig 5). Interestingly, DPP-4 i treatment mainly inhibited influx of M1 subset (*P* < 0.02) along with an upward trend of M2 infiltration from the total macrophages (Fig 6C, 6D, and E). This suggests that DPP-4 i diminished total monocyte/macrophages accumulation in the ischemia-injured tissue, and converted them into the M2 population, which displays strong anti-inflammatory, reparative and angiogenic functions. Interestingly, the influx of stem or progenitor cell (CD117 and Sca-1) was greater in DIO group (*P* < 0.02) than in the MK-0226-treated DIO group (Fig 6F and 6G).

**Fig 5.**
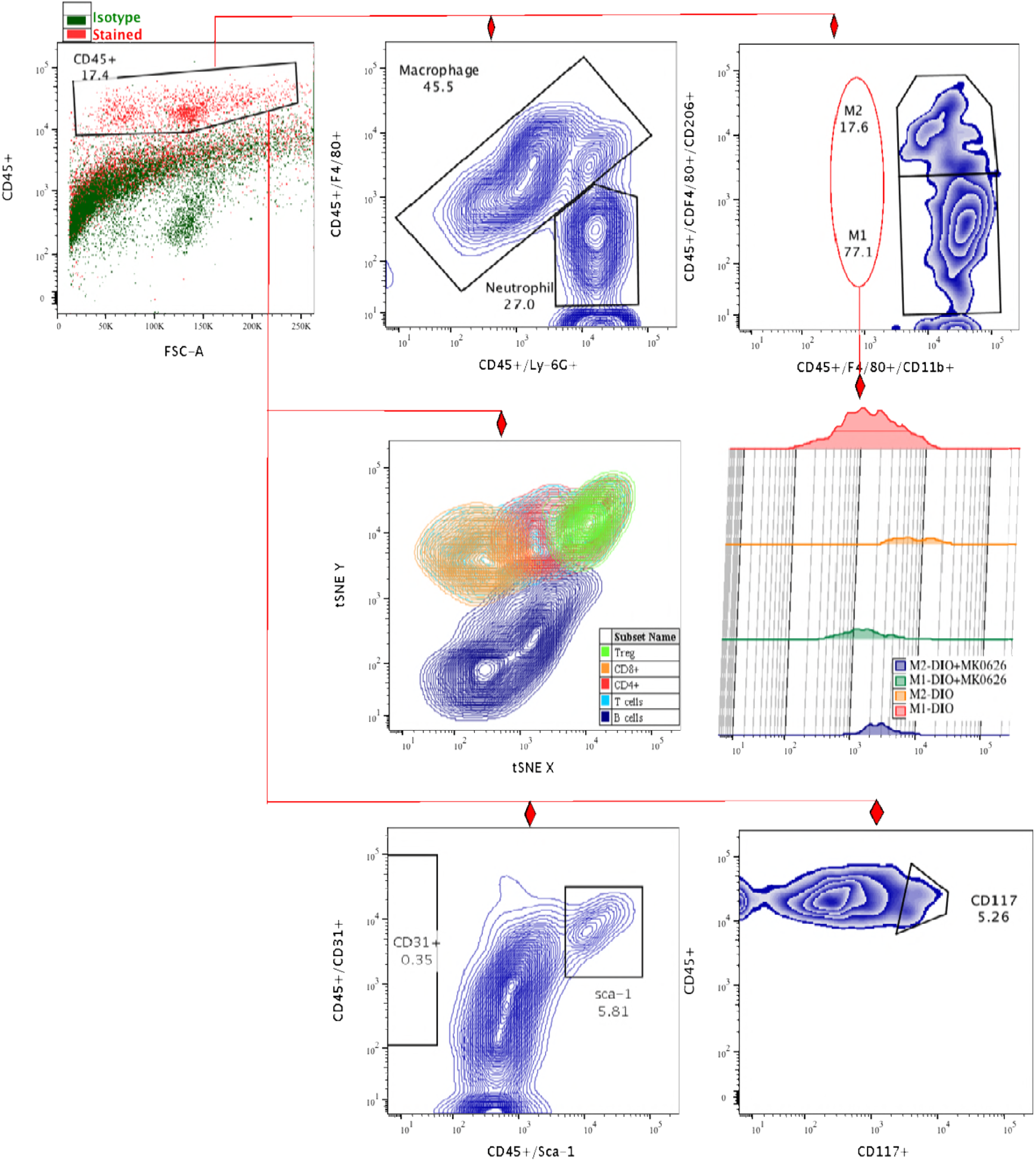
Flow cytometric gating strategy of ischemic tissue infiltrated cells. The numbers of recruited total T-cells (CD45^+^/CD3e^+^), subsets of T-helper cells (CD45^+^/CD3e^+^/CD4^+^, *P* < 0.02), and cytotoxic T-cells (CD45^+^/CD3^+^/CD8a^+^, P = 0.2) significantly decreased, while the frequency of regulatory T-cells (CD45^+^/CD4^+^/CD25^+^) tended to increase in the MK-0626-administered DIO group, in comparison with DIO group (Figs 6H–K). In tSNE analysis, B-cell (CD19^+^) recruitment into the LHI tissue showed an upward trend in the DIO group, in comparison with the MK0626-administered DIO group, although this difference was not statistically significant (*P* > 0.11) (Fig 6L)

**Fig 6.**
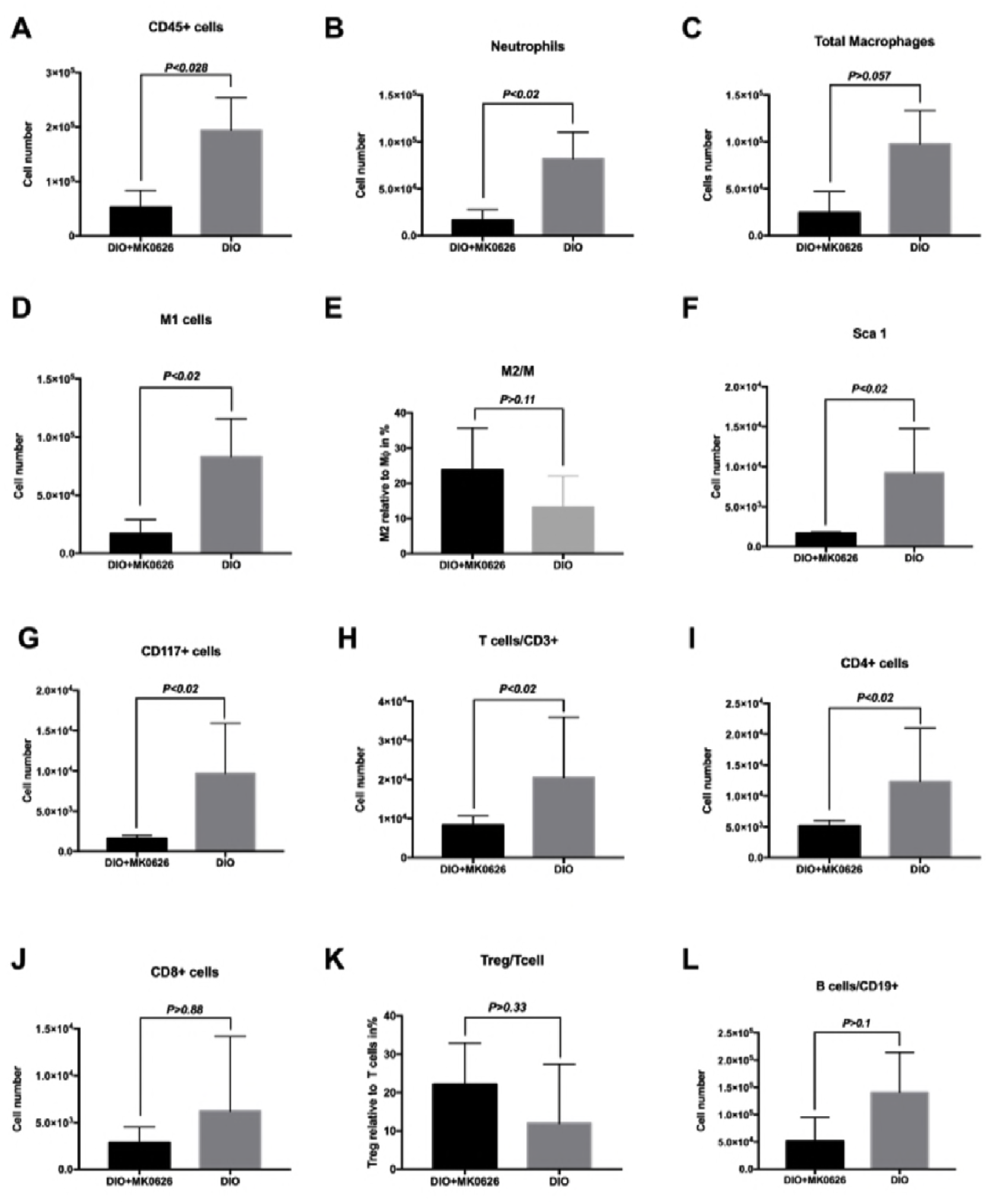
Quantification and characterization of ischemic tissue-infiltrated cells. (A–L, except E and K) All values are the absolute number of cells that infiltrated per anterior tibial muscle (n = 4–5, each). (E) M2 to total macrophages and (K) Relative ratio of T_reg_ cells to total T cells in CD45^+^ cells (n = 4–5, each). Data were analyzed by Mann-Whitney U test.

In summary, MK-0626 administration suppressed the influx of all hematopoietic lineage cells (mainly pro-inflammatory cells) and accelerated regeneration-associated cell (RACs) polarization from pro-inflammatory M1 and CD4^+^ cells to the anti-inflammatory/reparative M2 and T_reg_ cells, thus inhibiting unproductive inflammatory cascades following ischemic injury for further beneficial tissue restoration.

### MK-0626 administration enhanced MVD and pericyte recruitment in ischemic tissues

To evaluate MVD and pericyte recruited arterioles, IsolectineB4 and α-SMAI co-staining was performed. The immunohistology study revealed that in MK-0626-treated animals MVD was dramatically increased in comparison with DIO control (MVD/mm^2^; 872 ± 56 in MK-0626-treated vs. 528 ± 44 in DIO control, *P* < 0.001)(Fig 7A, a–d and 7B). To verify effect of MK-0626-treatment on arterial maturation, we stained ischemic hind limb tissues with α-SMA to detect pericyte recruitment. As shown in Fig 7A(e–h) and 7C, the number of αSMA positive vessels were superior in MK-0226-administered DIO mice (204 ± 81, P < 0.05), compared with that in DIO mice (160 ± 94). These results indicate that MK-0626 administration promoted MVD enhancement along with pericyte recruitment for vascular maturation after ischemic injury.

**Fig 7.**
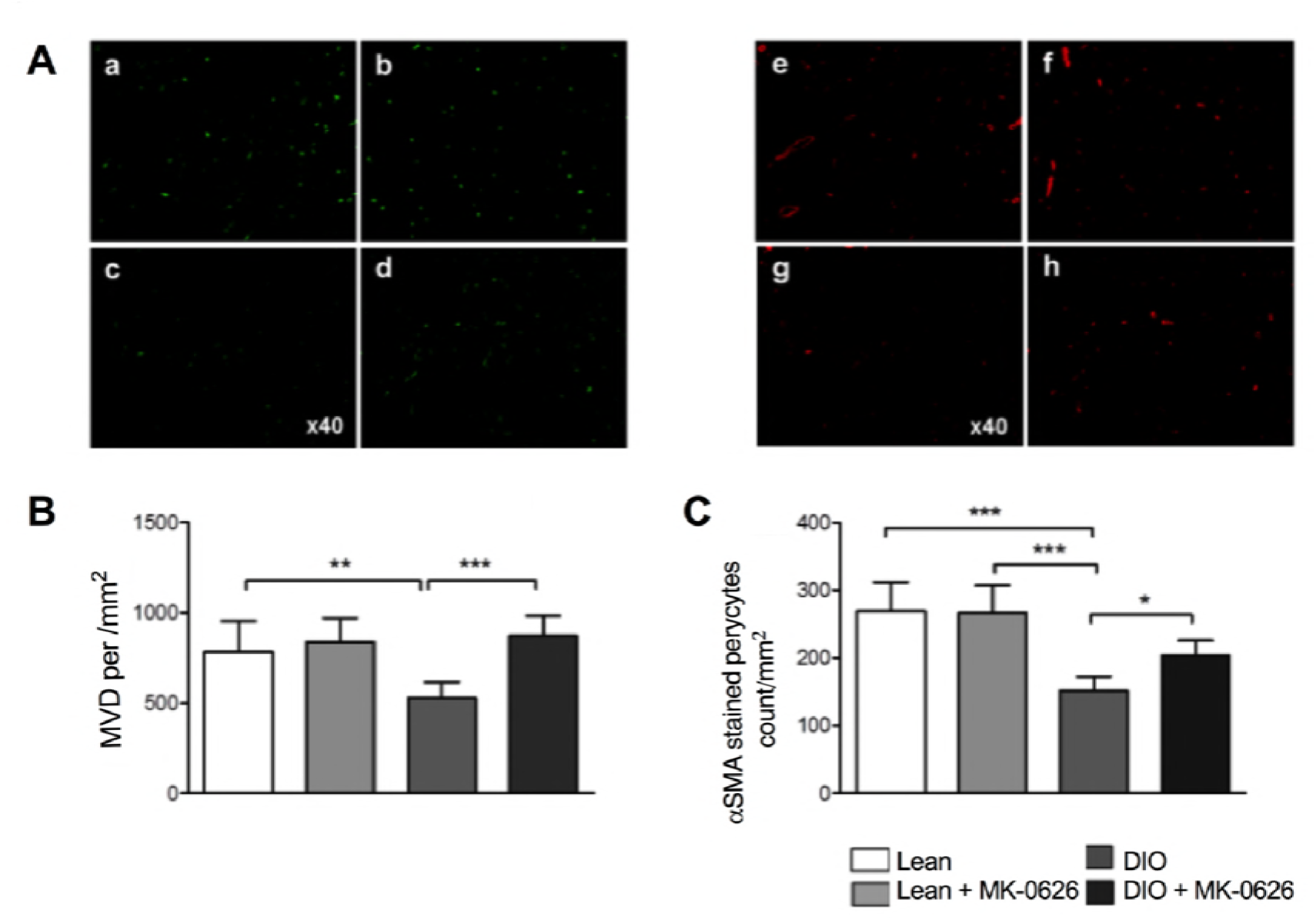
MK-0626-treatment promotes vascular regeneration in ischemic hind limb. (A) Representative pictures of angiogenesis and arteriogenesis in anterior tibial muscle (ATM) in each group. (a–d) The mouse microvessels were stained with FITCconjugated isolectin B4 (green). (e–h) Pericyte-recruited microvessels were stained with Cy3-conjugated anti-a SMA antibody (red). (B) The microvessels were counted with FITC-conjugated isolectin B4 under the HPF of 40× on a fluorescence microscope (C). The pericyte-recruited microvessels were counted with Cy3-conjugated anti-α-SMA antibody. Data are represented as mean ± SE. N = 6 mice per group. Experiments were repeated twice with similar results. In the graph, \**P* < 0.05, \*\**P* < 0.01, and \*\*\**P* < 0.001 as determined by One-Way ANOVA and Dunn’s multiple comparison test.

## Discussion

In the present study, we have demonstrated that the DPP-4 inhibitor MK-0626 enhanced vascular development by promoting EPC differentiation and orchestrating regenerative microenvironment of ischemic tissues through reduced influx of pro-inflammatory cells, such as neutrophils, M1 macrophages, cytotoxic T-cells and B-cells, and instead recruiting regeneration-associated cells, such as M2 macrophages and T_reg_ cells in DIO mice following onset of acute HLI (Fig 8).

**Fig 8.**
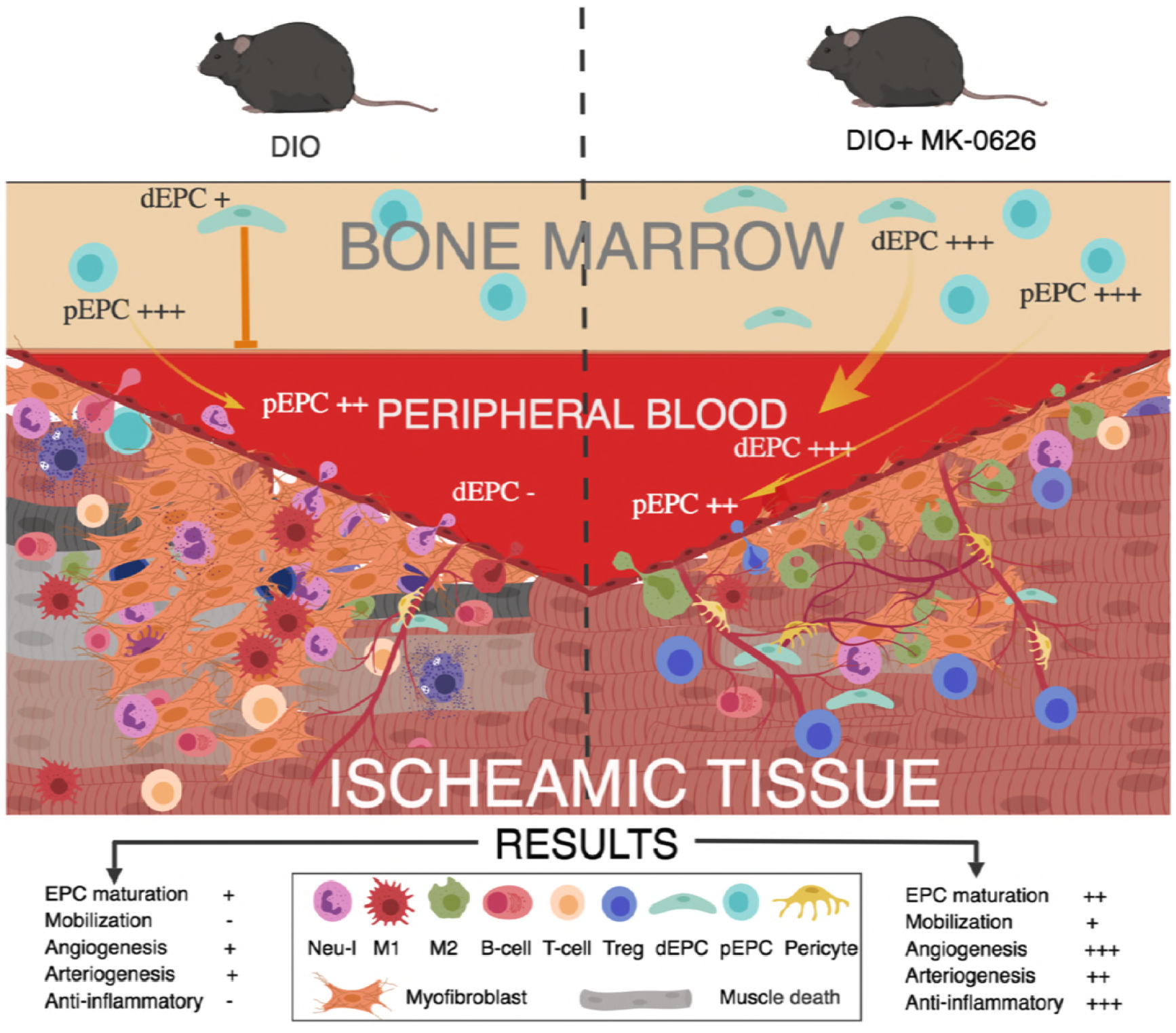
Summary of our study.

### Effect of MK-0626 on EPC bioactivity

First, we examined EPC colony forming abilities in PB of animals to evaluate MK-0626-triggered EPC mobilization effects. Definitive EPCs (dEPCs), representing more differentiated colony forming EPCs than primitive EPCs (pEPCs), significantly decreased in PB of DIO mice compared with healthy Lean mice. This impairment of dEPC kinetics in DIO was abrogated upon MK-0626 administration. Further, when we performed the same assay on BM-EPCs, represented by KSL cells, we found no differences between the groups of DIO and lean mice, suggesting that the impairment of dEPC kinetics in PB may be based on the obese condition. Moreover, the calculated number of EPC-CFUs in hemi bone displayed stimulated expansion of EPCs in BM microenvironment in DIO mice, and MK-0626 did not affect the frequency of EPC-CFUs in BM in both DIO and Lean mice. Taken together, our data suggest that the obese condition decreased EPC kinetics in PB through impairment in EPC differentiation, but stimulated EPC expansion in BM microenvironment. Further, MK-0626 recovered the impaired dEPC kinetics into circulation, possibly by mobilization effect, but could not stimulate further EPC expansion in BM microenvironment.

Former experiments showed that DIO condition aggravated stromal derived factor-1a (SDF-1a, CXCL12/CXCR4) axis, which is crucial for guiding hematopoietic cells, including EPCs, to the site of injury. SDF-1 is a substrate of DPP-4, and MK-0626 suppress degradation of SDF-1 by inhibiting enzyme activity of DPP-IV, consequently, the SDF-1 increases biological activities in PB and BM [10]. In accordance with this, our finding indicated that diabetic condition specifically impaired function of dEPC mobilization from BM to circulation. Under such conditions, DPP-4 i might restore the mobilization through preservation of SDF-1 protein by modulating enzyme activity of CD26/DPP-IV.

### MK-0626 effect on vascular development

In vivo transplantation experiment demonstrated that while limb perfusion recovery of DIO animal at day 14 was significantly decreased in comparison with healthy Lean mice recovery, MK-0626 treatment abrogated the deterioration in DIO ischemic mice. Immunohistology analysis revealed that vascular regeneration, such as IB4 stained capillary densities as well as pericyte-triggered arteriole maturation, were superior in MK-0626-administered DIO group, likely because the DPP-4 i promotes differentiation of primitive EPC to the definitive EPC to stimulate angiogenesis and arteriogenesis. In acute arterial injury model, short-term inhibition of DPP-4 i enhances endothelial regeneration through inhibition of SDF-1a degradation, and consequently increasing recruitment of circulating endothelial progenitor cells crucial for blood vessel development [20]. DPP-4 i is considered to promote vascular regeneration by two synergistic effects: by inhibiting the degradation of SDF-1a and via anti-inflammatory effects. Such incretin-based therapies with DP-4IV i, displays anti-inflammatory activity through glucose control via DPP-4 [21] Pathological and physiological angiogenesis initiation mainly occur after inflammation [22]. However, on T2DM, chronic inflammation induces excessive inflammatory cytokine secretion, which affects stem cell and EPC differentiation for effective vascular regeneration [23].

### Anti-inflammatory effect of MK-0626 to the ischemic tissue

DPP-4 i has been reported as an anti-inflammatory medicine in chronic inflammatory diseases, including T2DM [24],[25]. We assumed this DPP-4 i anti-inflammatory effect might contribute to vascularization and tissue regeneration, especially in situ ischemic tissue through recruitment of immune cells [26, 27]. Our hematopoietic cell isolation experiment from ischemic muscles suggested that DPP-4 i had an inhibitory effect on unproductive inflammation by decreasing influx of total blood cell accumulation (by 5-fold), and pro-inflammatory cells such as neutrophils (by 6.2-fold), total macrophages (by 7.4-fold), M1 (by 7-fold), total T-cells (by 2.2-fold), cytotoxic T-cells (by 1.6-fold), and B-cells (by 4-fold), and by increasing regeneration-associated cells, such as M2/M ratio (by 2-fold) and T_reg_/T-helper ratio (by 2-fold). Recent research also highlighted that DPP-4 inhibition attenuates obesity-related inflammation, atherosclerosis, and insulin resistance by regulating M1/M2 macrophage polarization [28, 29]. In an *<u>ApoE</u>*^−/−^ mouse model on high cholesterol diet, long-term treatment with the DPP4 inhibitor sitaglipin significantly reduced atherosclerotic plaque, and this effect was inversely correlated with number of M2 macrophages in the plaque. Blockade of CXCR4/SDF-1 signaling by AMD3100 inhibited aortic M2 accumulation and the therapeutic effect of Sitagliptin [28]. Interestingly, in our study, c-Kit and Sca-1 cells significantly infiltrated into the DIO mice ischemic tissues, in comparison with the MK-0626-treated counterpart. Our ex vivo data revealed that proportion of primitive EPCs was higher than definitive EPCs in PB and BM in DIO mice, indicating that obesity-induced inflammatory milieu decrease stem/progenitor cell differentiation in ischemic tissue due to unproductive inflammation, which supports other studies [23, 30, 31]. The DPP-4 inhibitors, such as Sitagliptin [32], Vildagliptin [33], Linagliptin [34], Teneligliptin [35], Anagliptin [36], Trelagliptin [37], and Omarigliptin [38] are already in use for clinical treatment of diabetes. Sitagliptin increases the number of circulating progenitor cells in mouse models [39] and T2DM patients [10, 40]. Similar to Sitagliptin, in our study, MK-0626 also increased the number of circulating endothelial progenitor cells in healthy mouse. Many research groups have reported that to increase the number of circulating EPCs in T2DM, EPCs kinetics and the vascular regeneration is important to prevent the disease [41, 42]. Some have also reported that DPP-4 i decreases the risk of cardiovascular disease in T2DM [43–46].

To conclude, our study shows that DPP-4 inhibition has a beneficial effect on vasculogenesis by enhancing EPC differentiation and bioactivity. Moreover, DPP-4 inhibition decreased the recruitment of pro-inflammatory cells into the ischemic injury, along with an increase in regeneration-associated cells, the latter being important in the tissue restoration and regeneration. Further clinical trials on metabolic syndrome need be conducted to prove the therapeutic potential of DPP-4 i in clinical practice as preventive medicine of arteriosclerosis.

## Acknowledgments

We thank to the Support Center for Medical Research and Education, Tokai University School of Medicine for excellent technical assistance and animal care.

## References

1. Asahara T, Murohara T, Sullivan A, Silver M, van der Zee R, Li T, et al. Isolation of putative progenitor endothelial cells for angiogenesis. Science. 1997;275(5302):964–7. Epub 1997/02/14. PubMed PMID: 9020076.

2. Takahashi T, Kalka C, Masuda H, Chen D, Silver M, Kearney M, et al. Ischemia-and cytokine-induced mobilization of bone marrow-derived endothelial progenitor cells for neovascularization. Nature medicine. 1999;5(4):434–8. Epub 1999/04/15. doi: 10.1038/7434. PubMed PMID: 10202935.

3. Tepper OM, Galiano RD, Capla JM, Kalka C, Gagne PJ, Jacobowitz GR, et al. Human endothelial progenitor cells from type II diabetics exhibit impaired proliferation, adhesion, and incorporation into vascular structures. Circulation. 2002;106(22):2781–6. Epub 2002/11/27. PubMed PMID: 12451003.

4. Fadini GP, Ciciliot S, Albiero M. Concise Review: Perspectives and Clinical Implications of Bone Marrow and Circulating Stem Cell Defects in Diabetes. Stem Cells. 2017;35(1):106–16. Epub 2016/07/13. doi: 10.1002/stem.2445. PubMed PMID: 27401837.

5. Yamaguchi J, Kusano KF, Masuo O, Kawamoto A, Silver M, Murasawa S, et al. Stromal cell-derived factor-1 effects on ex vivo expanded endothelial progenitor cell recruitment for ischemic neovascularization. Circulation. 2003;107(9):1322–8. Epub 2003/03/12. PubMed PMID: 12628955.

6. Matsumoto T, Ii M, Nishimura H, Shoji T, Mifune Y, Kawamoto A, et al. Lnk-dependent axis of SCF-cKit signal for osteogenesis in bone fracture healing. The Journal of experimental medicine. 2010;207(10):2207–23. Epub 2010/09/22. doi: 10.1084/jem.20100321. PubMed PMID: 20855498; PubMed Central PMCID: PMC2947078.

7. Deacon CF. Peptide degradation and the role of DPP-4 inhibitors in the treatment of type 2 diabetes. Peptides. 2018;100:150–7. doi: https://doi.org/10.1016/j.peptides.2017.10.011.

8. Jungraithmayr W, De Meester I, Matheeussen V, Baerts L, Arni S, Weder W. CD26/DPP-4 inhibition recruits regenerative stem cells via stromal cell-derived factor-1 and beneficially influences ischaemia-reperfusion injury in mouse lung transplantation. European journal of cardio-thoracic surgery: official journal of the European Association for Cardio-thoracic Surgery. 2012;41(5):1166–73. Epub 2012/01/06. doi: 10.1093/ejcts/ezr180. PubMed PMID: 22219460.

9. Wang CH, Cherng WJ, Yang NI, Hsu CM, Yeh CH, Lan YJ, et al. Cyclosporine increases ischemia-induced endothelial progenitor cell mobilization through manipulation of the CD26 system. American journal of physiology Regulatory, integrative and comparative physiology. 2008;294(3):R811–8. Epub 2007/12/21. doi: 10.1152/ajpregu.00543.2007. PubMed PMID: 18094068.

10. Fadini GP, Boscaro E, Albiero M, Menegazzo L, Frison V, de Kreutzenberg S, et al. The oral dipeptidyl peptidase-4 inhibitor sitagliptin increases circulating endothelial progenitor cells in patients with type 2 diabetes: possible role of stromal-derived factor-1alpha. Diabetes care. 2010;33(7):1607–9. Epub 2010/04/02. doi: 10.2337/dc10-0187. PubMed PMID: 20357375; PubMed Central PMCID: PMC2890368.

11. Masuda H, Alev C, Akimaru H, Ito R, Shizuno T, Kobori M, et al. Methodological development of a clonogenic assay to determine endothelial progenitor cell potential. Circ Res. 2011;109(1):20–37. Epub 2011/05/14. doi: 10.1161/circresaha.110.231837. PubMed PMID: 21566217.

12. Edmondson SD, Mastracchio A, Mathvink RJ, He J, Harper B, Park YJ, et al. (2S,3S)-3-Amino-4-(3,3-difluoropyrrolidin-1-yl)-N,N-dimethyl-4-oxo-2-(4-[1,2,4]tr iazolo[1,5-a]-pyridin-6-ylphenyl)butanamide: a selective alpha-amino amide dipeptidyl peptidase IV inhibitor for the treatment of type 2 diabetes. Journal of medicinal chemistry. 2006;49(12):3614–27. Epub 2006/06/09. doi: 10.1021/jm060015t. PubMed PMID: 16759103.

13. Aroor AR, Habibi J, Ford DA, Nistala R, Lastra G, Manrique C, et al. Dipeptidyl peptidase-4 inhibition ameliorates Western diet-induced hepatic steatosis and insulin resistance through hepatic lipid remodeling and modulation of hepatic mitochondrial function. Diabetes. 2015;64(6):1988–2001. Epub 2015/01/22. doi: 10.2337/db14-0804. PubMed PMID: 25605806; PubMed Central PMCID: PMCPMC4439570.

14. Masuda H, Tanaka R, Fujimura S, Ishikawa M, Akimaru H, Shizuno T, et al. Vasculogenic Conditioning of Peripheral Blood Mononuclear Cells Promotes Endothelial Progenitor Cell Expansion and Phenotype Transition of Anti-Inflammatory Macrophage and T Lymphocyte to Cells With Regenerative Potential. Journal of the American Heart Association. 2014;3(3). doi: 10.1161/jaha.113.000743. PubMed PMID: WOS:000209562400020.

15. Asahara T, Masuda H, Takahashi T, Kalka C, Pastore C, Silver M, et al. Bone marrow origin of endothelial progenitor cells responsible for postnatal vasculogenesis in physiological and pathological neovascularization. Circ Res. 1999;85(3):221–8. Epub 1999/08/07. PubMed PMID: 10436164.

16. Masuda H, Tanaka R, Fujimura S, Ishikawa M, Akimaru H, Shizuno T, et al. Vasculogenic conditioning of peripheral blood mononuclear cells promotes endothelial progenitor cell expansion and phenotype transition of anti-inflammatory macrophage and T lymphocyte to cells with regenerative potential. Journal of the American Heart Association. 2014;3(3):e000743. Epub 2014/06/27. doi: 10.1161/JAHA.113.000743. PubMed PMID: 24965023.

17. Pinto AR, Chandran A, Rosenthal NA, Godwin JW. Isolation and analysis of single cells from the mouse heart. Journal of immunological methods. 2013;393(1-2):74–80. Epub 2013/04/13. doi: 10.1016/j.jim.2013.03.012. PubMed PMID: 23578979.

18. Lumeng CN, Saltiel AR. Inflammatory links between obesity and metabolic disease. J Clin Invest. 1212011. p. 2111–7.

19. Monteiro R, Azevedo I. Chronic Inflammation in Obesity and the Metabolic Syndrome. Mediators Inflamm. 2010;2010. doi: 10.1155/2010/289645. PubMed PMID: 20706689; PubMed Central PMCID: PMC2913796.

20. Brenner C, Kränkel N, Kühlenthal S, Israel L, Remm F, Fischer C, et al. Short-term inhibition of DPP-4 enhances endothelial regeneration after acute arterial injury via enhanced recruitment of circulating progenitor cells. International Journal of Cardiology. 2014;177(1):266–75. doi: https://doi.org/10.1016/j.ijcard.2014.09.016.

21. Scheen AJ, Esser N, Paquot N. Antidiabetic agents: Potential anti-inflammatory activity beyond glucose control. Diabetes & metabolism. 2015;41(3):183–94. Epub 2015/03/22. doi: 10.1016/j.diabet.2015.02.003. PubMed PMID: 25794703.

22. Carmeliet P. Angiogenesis in life, disease and medicine. Nature. 2005;438(7070):932–6. Epub 2005/12/16. doi: 10.1038/nature04478. PubMed PMID: 16355210.

23. Desouza CV, Hamel FG, Bidasee K, O’Connell K. Role of inflammation and insulin resistance in endothelial progenitor cell dysfunction. Diabetes. 2011;60(4):1286–94. Epub 2011/02/25. doi: 10.2337/db10-0875. PubMed PMID: 21346178; PubMed Central PMCID: PMC3064102.

24. Yazbeck R, Howarth GS, Abbott CA. Dipeptidyl peptidase inhibitors, an emerging drug class for inflammatory disease? Trends in pharmacological sciences. 2009;30(11):600–7. Epub 2009/10/20. doi: 10.1016/j.tips.2009.08.003. PubMed PMID: 19837468.

25. Satoh-Asahara N, Sasaki Y, Wada H, Tochiya M, Iguchi A, Nakagawachi R, et al. A dipeptidyl peptidase-4 inhibitor, sitagliptin, exerts anti-inflammatory effects in type 2 diabetic patients. Metabolism: clinical and experimental. 2013;62(3):347–51. Epub 2012/10/16. doi: 10.1016/j.metabol.2012.09.004. PubMed PMID: 23062489.

26. Aliotta JM, Pereira M, Wen S, Dooner MS, Del Tatto M, Papa E, et al. Bone Marrow Endothelial Progenitor Cells Are the Cellular Mediators of Pulmonary Hypertension in the Murine Monocrotaline Injury Model. Stem Cells Transl Med. 2017;6(7):1595–606. Epub 2017/05/06. doi: 10.1002/sctm.16-0386. PubMed PMID: 28474513; PubMed Central PMCID: PMCPMC5689760.

27. Zhao Y, Yang L, Wang X, Zhou Z. The new insights from DPP-4 inhibitors: their potential immune modulatory function in autoimmune diabetes. Diabetes/metabolism research and reviews. 2014;30(8):646–53. Epub 2014/01/22. doi: 10.1002/dmrr.2530. PubMed PMID: 24446278.

28. Brenner C, Franz WM, Kühlenthal S, Kuschnerus K, Remm F, Gross L, et al. DPP-4 inhibition ameliorates atherosclerosis by priming monocytes into M2 macrophages. International Journal of Cardiology. 2015;199:163–9. doi: https://doi.org/10.1016/j.ijcard.2015.07.044.

29. Zhuge F, Ni Y, Nagashimada M, Nagata N, Xu L, Mukaida N, et al. DPP-4 inhibition by linagliptin attenuates obesity-related inflammation and insulin resistance by regulating M1/M2 macrophage polarization. Diabetes. 2016;db160317. doi: 10.2337/db16-0317.

30. Avogaro A, Albiero M, Menegazzo L, Kreutzenberg Sd, Fadini GP. Endothelial Dysfunction in Diabetes. Diabetes. 2011;34. doi: 10.2337/dc11-s239.

31. Matsubara J, Sugiyama S, Akiyama E, Iwashita S, Kurokawa H, Ohba K, et al. Dipeptidyl Peptidase-4 Inhibitor, Sitagliptin, Improves Endothelial Dysfunction in Association With Its Anti-Inflammatory Effects in Patients With Coronary Artery Disease and Uncontrolled Diabetes. Circulation journal: official journal of the Japanese Circulation Society. 2013. Epub 2013/02/07. PubMed PMID: 23386232.

32. Shimoda S, Iwashita S, Ichimori S, Matsuo Y, Goto R, Maeda T, et al. Efficacy and safety of sitagliptin as add-on therapy on glycemic control and blood glucose fluctuation in Japanese type 2 diabetes subjects ongoing with multiple daily insulin injections therapy. Endocrine journal. 2013;60(10):1207–14. Epub 2013/08/06. PubMed PMID: 23912974.

33. Russo E, Penno G, Del Prato S. Managing diabetic patients with moderate or severe renal impairment using DPP-4 inhibitors: focus on vildagliptin. Diabetes, metabolic syndrome and obesity: targets and therapy. 2013;6:161–70. Epub 2013/05/08. doi: 10.2147/DMSO.S28951. PubMed PMID: 23650450; PubMed Central PMCID: PMC3639752.

34. Hoimark L, Laursen T, Rungby J. Potential role of linagliptin as an oral once-daily treatment for patients with type 2 diabetes. Diabetes, metabolic syndrome and obesity: targets and therapy. 2012;5:295–302. Epub 2012/09/07. doi: 10.2147/DMSO.S16288. PubMed PMID: 22952411; PubMed Central PMCID: PMC3430084.

35. Kishimoto M. Teneligliptin: a DPP-4 inhibitor for the treatment of type 2 diabetes. Diabetes, metabolic syndrome and obesity: targets and therapy. 2013;6:187–95. Epub 2013/05/15. doi: 10.2147/DMSO.S35682. PubMed PMID: 23671395; PubMed Central PMCID: PMC3650886.

36. Nishio S, Abe M, Ito H. Anagliptin in the treatment of type 2 diabetes: safety, efficacy, and patient acceptability. Diabetes, metabolic syndrome and obesity: targets and therapy. 2015;8:163–71. Epub 2015/04/04. doi: 10.2147/DMSO.S54679. PubMed PMID: 25834461; PubMed Central PMCID: PMC4370682.

37. Inagaki N, Onouchi H, Maezawa H, Kuroda S, Kaku K. Once-weekly trelagliptin versus daily alogliptin in Japanese patients with type 2 diabetes: a randomised, double-blind, phase 3, non-inferiority study. The lancet Diabetes & endocrinology. 2015;3(3):191–7. Epub 2015/01/23. doi: 10.1016/S2213-8587(14)70251-7. PubMed PMID: 25609193.

38. Sheu WH, Gantz I, Chen M, Suryawanshi S, Mirza A, Goldstein BJ, et al. Safety and Efficacy of Omarigliptin (MK-3102), a Novel Once-Weekly DPP-4 Inhibitor for the Treatment of Patients With Type 2 Diabetes. Diabetes care. 2015;38(11):2106–14. Epub 2015/08/28. doi: 10.2337/dc15-0109. PubMed PMID: 26310692.

39. Huang CY, Shih CM, Tsao NW, Lin YW, Huang PH, Wu SC, et al. Dipeptidyl peptidase-4 inhibitor improves neovascularization by increasing circulating endothelial progenitor cells. British journal of pharmacology. 2012;167(7):1506–19. Epub 2012/07/14. doi: 10.1111/j.1476-5381.2012.02102.x. PubMed PMID: 22788747; PubMed Central PMCID: PMC3514763.

40. Aso Y, Jojima T, Iijima T, Suzuki K, Terasawa T, Fukushima M, et al. Sitagliptin, a dipeptidyl peptidase-4 inhibitor, increases the number of circulating CD34(+)CXCR4(+) cells in patients with type 2 diabetes. Endocrine. 2015;50(3):659–64. Epub 2015/07/26. doi: 10.1007/s12020-015-0688-5. PubMed PMID: 26209038.

41. Cicek FA, Tokcaer-Keskin Z, Ozcinar E, Bozkus Y, Akcali KC, Turan B. Di-peptidyl peptidase-4 inhibitor sitagliptin protects vascular function in metabolic syndrome: possible role of epigenetic regulation. Molecular biology reports. 2014;41(8):4853–63. Epub 2014/05/20. doi: 10.1007/s11033-014-3392-2. PubMed PMID: 24838371.

42. Renna NF, Diez EA, Miatello RM. Effects of dipeptidyl-peptidase 4 inhibitor about vascular inflammation in a metabolic syndrome model. PloS one. 2014;9(9):e106563. Epub 2014/09/04. doi: 10.1371/journal.pone.0106563. PubMed PMID: 25184237; PubMed Central PMCID: PMC4153656.

43. Fadini GP, Avogaro A. Cardiovascular effects of DPP-4 inhibition: beyond GLP-1. Vascular pharmacology. 2011;55(1-3):10–6. Epub 2011/06/15. doi: 10.1016/j.vph.2011.05.001. PubMed PMID: 21664294.

44. Connelly KA, Zhang Y, Advani A, Advani SL, Thai K, Yuen DA, et al. DPP-4 inhibition attenuates cardiac dysfunction and adverse remodeling following myocardial infarction in rats with experimental diabetes. Cardiovascular therapeutics. 2013;31(5):259–67. Epub 2012/09/12. doi: 10.1111/1755-5922.12005. PubMed PMID: 22963483.

45. Yousefzadeh P, Wang X. The effects of dipeptidyl peptidase-4 inhibitors on cardiovascular disease risks in type 2 diabetes mellitus. Journal of diabetes research. 2013;2013:459821. Epub 2013/05/28. doi: 10.1155/2013/459821. PubMed PMID: 23710467; PubMed Central PMCID: PMC3654348.

46. Scheen AJ. Cardiovascular effects of DPP-4 inhibitors: from risk factors to clinical outcomes. Postgraduate Medicine. 2016.

